# The zebrafish (*Danio rerio*) snoRNAome

**DOI:** 10.1101/2024.07.26.605292

**Authors:** Renáta Hamar, Máté Varga

## Abstract

Small nucleolar RNAs (snoRNAs) are one of the most abundant and evolutionary ancient group of functional non-coding RNAs. They were originally described as guides of post-transcriptional rRNA modifications, but emerging evidence suggests that snoRNAs fulfil an impressive variety of cellular functions, some of which are only being discovered. To reveal the true complexity of snoRNA-dependent functions, we need to catalogue first the complete repertoire of snoRNAs in a given cellular context. While the systematic mapping and characterization of “snoRNAomes” for some species have been described recently, this has not been done hitherto for the zebrafish (*Danio rerio*). Using size-fractionated RNA sequencing data from adult zebrafish tissues, we created an interactive “snoRNAome” database for this species. Our custom-designed analysis pipeline allowed us to identify with high-confidence 68 previously unannotated snoRNAs in the zebrafish genome, resulting in the most complete set of snoRNAs to date in this species. Reanalyzing multiple previously published datasets, we also provide evidence for the dynamic expression of some snoRNAs during the early stages of zebrafish development and tissue-specific expression patterns for others in adults. To facilitate further investigations into the functions of snoRNAs in zebrafish, we created a novel interactive database, snoDanio, which can be used to explore small RNA expression from transcriptomic data.

## INTRODUCTION

Small nucleolar RNAs (snoRNAs) form one of the most abundant and ancient group of functional non-coding RNAs (ncRNAs). Their main role is to guide the chemical modification of specific nucleosides in several RNA classes. Consequently, they play critical roles in multiple cellular regulatory processes, such as the maturation and nucleolytic processing of ribosomal RNAs (rRNAs), the chromatin architecture and alternative splicing (Han et al. 2022; Watkins and Bohnsack 2011; Khosraviani et al. 2019).

Based on common sequence motifs and conserved structural features snoRNAs are classified in two major families, C/D box and H/ACA box snoRNAs (Figure 1A, B), abbreviated as SNORDs and SNORAs, respectively (Bratkovič et al. 2019; Webster and Ghalei 2023). Members of both families are associated with larger ribonucleoprotein (RNP) complexes where they guide the function of partner enzymes to initiate the posttranscriptional modification of various RNA species. These enzymes, the methyltransferase fibrillarin (FBL) and the pseudouridine synthase dyskerin (DKC1), respectively, will catalyze the formation of ribose 2’-*O*-methyl (Nm) groups and the isomerization of uridine into pseudoridine (Ψ) (Garus and Autexier 2021; Shubina et al. 2018; Kiss et al. 2022).

**Figure 1:**
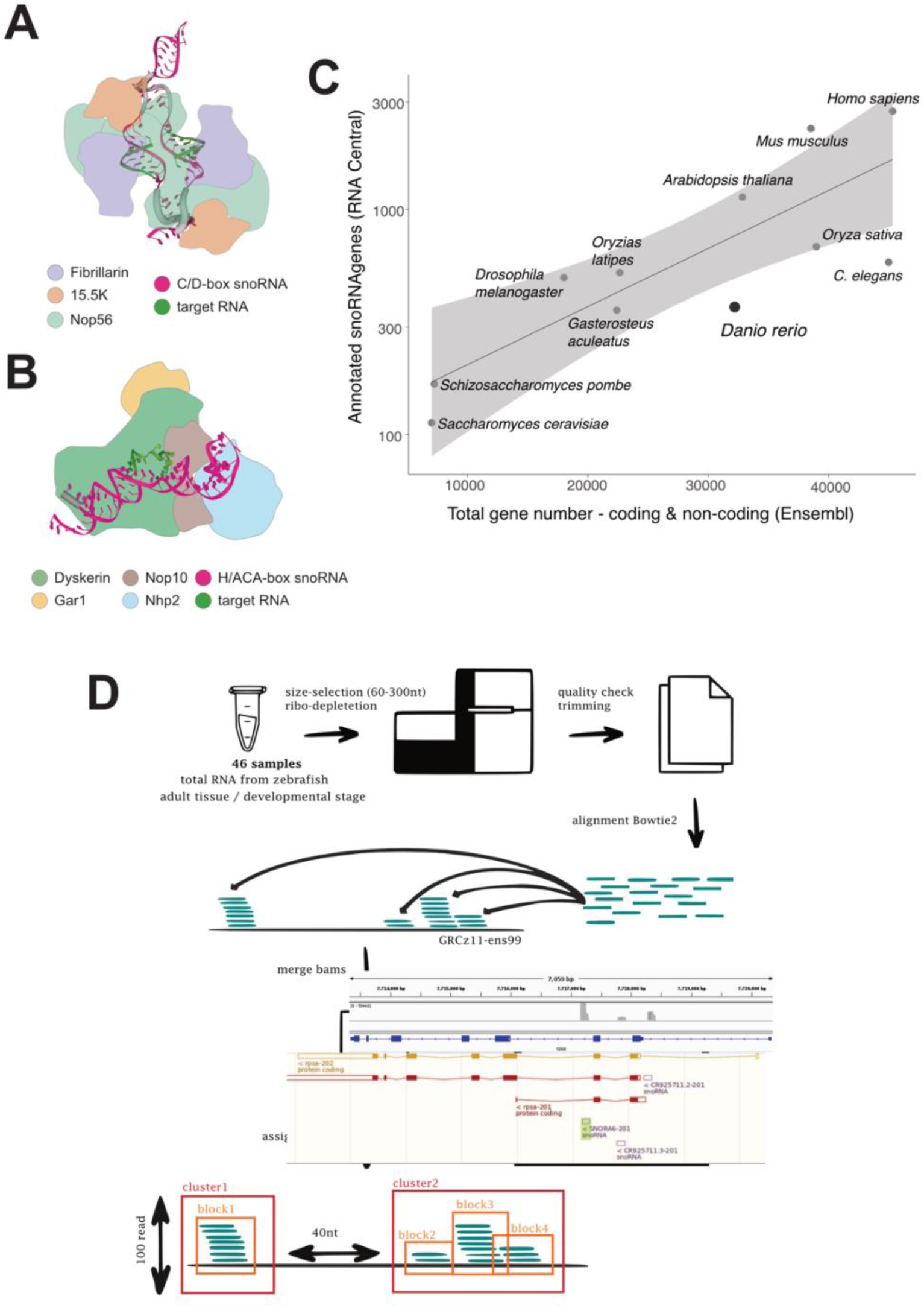
Annotation of novel snoRNAs in the zebrafish genome. (A,B) Different snoRNA types participate in different RNP complexes: whereas C/D box snoRNAs can be found in complex with Fibrillarin, Nop56 and 15.5K (A), H/ACA box snoRNAs form complexes with Dyskerin, Nop10, Gar1 and Nhp2 (B). (C) The ratio of annotated snoRNAs in the zebrafish genome is relatively low compared with other species. (D) The computational pipeline used to annotate novel snoRNAs from transcriptomic datasets.

While the predominant targets of SNORD-directed methylation appear to be rRNAs, with few snRNAs also modified by FBL (Gumienny et al. 2017), among the targets of DKC1 we can find snoRNAs, snRNAs, rRNAs and mRNAs as well (Karijolich et al. 2015). Both of the discussed post-transcriptional modifications seem to be essential for nucleolytic processing during rRNA maturation and in their absence ribosomal biogenesis is disrupted (Watkins and Bohnsack 2011). Defects of DKC1 function can lead to monogenic diseases like Dyskeratosis congenita (DC) (Savage 2022), Bowen-Conradi syndrome (BCS) (Armistead et al. 2009) or severe nephrotic syndrome with cataract (Balogh et al. 2020). Defective pseudouridylation of specific rRNA sites can also lead to cancer through the disruption of ribosomal function (McMahon et al. 2019, 2015; Liang et al. 2019; Huang et al. 2019; Babaian et al. 2020). Erroneous fibrillarin expression has also been linked to cancer (Marcel et al. 2013; Zhang et al. 2021; Long et al. 2022) and some SNORDs have been suggested to act either as oncogenes or as tumor suppressors (Zhu et al. 2019; Chen et al. 2015).

Emerging experimental evidence also suggests non-canonical functions for snoRNAs, linked to the genesis of other ncRNAs, mRNA 3’ processing, protein trapping and exosome recruitment to target RNAs (Huang et al. 2017; Bratkovič et al. 2019; Falaleeva and Stamm 2012; Bergeron et al. 2020).

Due to this large body of evidence about the multitude of crucial biological processes affected by snoRNAs, there have been several concerted efforts to map the complete snoRNA repertoire (so called snoRNAome) in several species (Canzler et al. 2017; Jorjani et al. 2016; Fafard-Couture et al. 2021).

Zebrafish rose to prominence as a developmental model system in the 1990s thanks to its numerous advantages (Varga 2018; Lieschke and Currie 2007). Aided by the dynamic expansion of the genetic toolbox it became one of the dominant models of biomedical research over the past couple of decades. This versatility also makes it an ideal subject to study the physiological roles of snoRNAs (Higa-Nakamine et al. 2012).

The relatively low number of known (annotated) zebrafish snoRNA genes (Figure 1C) suggests that the annotation of the snoRNA pool in this species is not yet complete, and that part of the zebrafish snoRNAome is still to be discovered. By reanalyzing previously published transcriptomic datasets and acquiring novel ones enriched in snoRNAs from multiple adult tissues we were able to identify and annotate several new snoRNAs. The analysis of the most complete zebrafish snoRNAome to date made it also possible to uncover the developmentally dynamic and/or tissue-specific expression of multiple snoRNAs in this species. We also present snoDanio (https://renata-h.shinyapps.io/98665a9405b44ede86eeb7179988104f/), an interactive database for the analysis of different transcriptomic datasets containing snoRNAs.

## MATERIALS AND METHODS

### Fish maintenance and RNA extraction

Adult wild-type fish used for these experiments were maintained in the animal facility of the Biology Institute of ELTE Eötvös Loránd University. Tissues were isolated from adult zebrafish based on standard protocols. All protocols employed in our study were approved by the Hungarian National Food Chain Safety Office (Permit XIV-I-001/515-4/2012). Two male and two female fish were sacrificed for isolation of total RNA from multiple tissues to produce four biological replicants. Extreme care was taken to avoid contamination and obtain pure homogenous tissue samples. The tissues were repeatedly washed in PBS to remove contaminating debris and TRI reagent (Zymo Research, Cat. No.: R2050) was used to collect total RNA, according to the manufacturer’s protocol.

### RNA library preparation and sequencing

All samples were subjected to size selection (60-200 bp) without ribodepletion before sequencing. Samples were sequenced by Novogene Ltd. on an Illumina NovaSeq 6000 PE150 platform. 20 million reads per sample were obtained through this strategy.

### Re-detection of previously annotated snoRNAs

We validated snoRNA prediction algorithms using the annotated version of the GRCz11 zebrafish genome (Ensembl version 104). After downloading the annotated snoRNA sequences, we applied three detection algorithms: cmsearch, snoReport, and snoscan/snoGPS with default parameters. The results of these searches were then combined to create a comprehensive list of re-detected, already described snoRNAs. This validation process aimed to enhance the reliability of snoRNA detection by cross-verifying predictions across multiple tools.

### snoRNA annotation pipeline

We created a novel, easily accessible and cloud-based pipeline, which greatly simplifies the identification of new snoRNA candidate sequences (Figure 1D). Our annotation pipeline is available on the Galaxy web platform (Community et al. 2024) via this link: https://usegalaxy.eu/u/danio/w/imported-annot-snos. Briefly, we used the splice-aware alignment algorithm HISAT2 to align reads to the zebrafish genome (GRCz11) and Bowtie2 to map reads in their continuum to the indexed reference. We considered searching for and filtering out further reads that map to rDNA sequences (Zhang et al. 2016), but since some snoRNAs are processed from rRNA, we decided against this approach. Bowtie2 mapped the reads to the genome in sensitive-local mode (Qu et al. 2015). We merged these files with the *blockcluster* algorithm (Bhatia et al. 2017), which also characterized the amount of reads. We set the filtering threshold to a minimum depth of 100 read per cluster and set the distance between clusters to 50 bp, as suggested in a previous study (Liu and Si 2014). In the consolidated file, we defined blocks from these reads and clusters from the blocks using BlockClust (Videm et al. 2014). We extracted the sections that overlapped with the annotations downloaded from ensembleV104. We used this approach for both time-series and tissue (both tRNA-seq) data. We merged the overlapping clusters by their genomic coordinates. For the 60-300 long ones, we downloaded the genomic sequence and ran it through tRNAscan-SE and the three known snoRNA scan applications to detect the corresponding sequences based on their structure. This method is good at detecting false positives, but it is worth analysing the transcriptome-matched portion of the sequencing to see if previously unknown snoRNA variants of genes described as pre-miR or lnc-RNA exist. Therefore, for these reads, we also need to look at the read profile of each gene to determine whether processed snoRNAs are present whose annotation is obscured by the parental non-coding gene. Based on the results of the RNAcentral sequence alignment run on predicted sequences we removed those containing sequences that were at least 95% identical to zebrafish non-coding sequences and could be identified as snoRNA (Supplementary Table 1).

### RNA-seq data processing pipelines for snoDanio

Our database includes RNA sequencing data from multiple BioProjects uploaded to the NCBI database (https://www.ncbi.nlm.nih.gov/bioproject/). As these projects contain raw RNA sequences obtained by different laboratories through different sequencing strategies, we created standardized pipelines for their processing. For consistency, we performed the alignment for each dataset starting from the raw data and mapped it to the most recent version (GRCz11) of the zebrafish genome. In each case the newly annotated snoRNAs were also added to the list of transcripts prior the mapping process, and we created novel Galaxy pipelines for the analysis of different RNA-seq experiment types (Table 1).

**Table 1:**
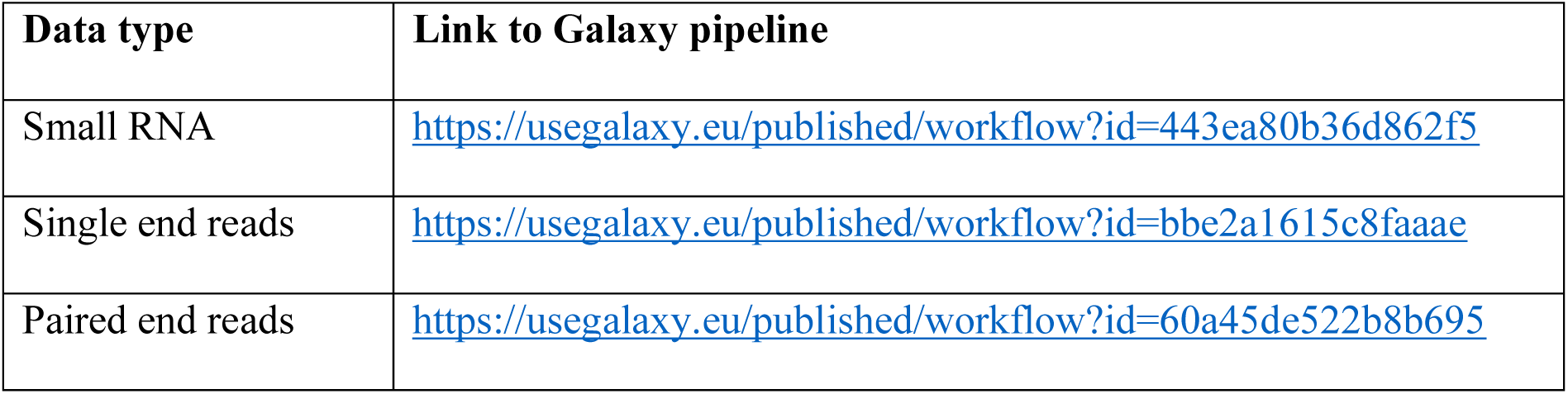
Different Galaxy pipelines used for the processing of external RNA-seq datasets.

| Data type | Link to Galaxy pipeline |
| --- | --- |
| Small RNA | https://usegalaxy.eu/published/workflow?id=443ea80b36d862f5 |
| Single end reads | https://usegalaxy.eu/published/workflow?id=bbe2a1615c8faaae |
| Paired end reads | https://usegalaxy.eu/published/workflow?id=60a45de522b8b695 |

To identify relevant BioProjects we downloaded the metadata for all zebrafish RNA sequencing experiments from the NCBI Sequence Read Archive (SRA) and searched for BioProjects that contain snoRNA size-range sequences. We filtered the results for the use of the HiSeq 2000 platform (as it matches the sequencing parameters of our own dataset) and excluded experimental designs that focus on polyA mRNA. The workflows resulted in new expression data for each experiment, in a columnar format. These datasets were collated to create R Data Serialization (rds) objects that form the core of our searchable snoDanio database (Hamar and Varga 2024).

### The snoDanio user interface

The snoDanio database pools previously available datasets with newly acquired sequencing data to create a comprehensive list of zebrafish snoRNAs, complemented with both snoRNA and host genes expression profiles in the examined datasets. The resulting database is accessible through a *shiny* web application (Chang et al. 2024) at https://renata-h.shinyapps.io/98665a9405b44ede86eeb7179988104f/.

The “Description of snoRNAs” section of our database allows the user to retrieve detailed information about various zebrafish snoRNA genes. For each gene the table showcases its ENSEMBL ID, source of annotation, and genomic location. By selecting a row in this table, one is able to see additional details about the chosen snoRNA in a dedicated sidebar panel. These additional informations include the gene symbol, the Rfam family, the parent gene information (when available), including the name and the biotype. We also provide the option to download the sequence of the selected snoRNA in FASTA format.

The “Explorer” part of the database is designed to provide an interactive platform for exploring the expression data of snoRNA genes across various BioProjects. Users can start by selecting a sequenced size range and an experiment type from the provided dropdown menus, which dynamically update the available project choices based on these selections. Upon choosing a project, the user can enter a specific snoRNA gene ID to investigate its expression levels. The database loads the rds object corresponding to the relevant BioProject and extracts the expression data for visualization. For the visualization of expression data the database generated plots using *ggplot2* and *plotly* (Wickham 2016; Sievert 2020). If the parent gene of the selected snoRNA is also available, a similar plot for the parent gene’s expression is generated. Users can also view a table listing the top five differentially expressed snoRNA genes for the selected project, facilitating quick identification of key genes of interest. This feature-rich interface enables comprehensive exploration and analysis of gene expression data, making it a valuable tool for researchers.

## RESULTS

### *De novo* annotation of zebrafish snoRNAs

The identification of snoRNAs is often difficult due to their lack of overall sequence conservation, their relatively small size (60-300 bp) and the shortness of the sequence motifs characteristic for this RNA class. It is not surprising, therefore, that multiple *in silico* methods based on different approaches have been developed for snoRNA prediction over the years. Some rely more heavily on purely structural features (*cmsearch*) (Cui et al. 2016), while others focus more on sequence similarities (*snoReport*) (Oliveira et al. 2016) and some mix the two approaches (*snoGPS, snoscan*) (Schattner et al. 2005).

To test the accuracy of these algorithms, we performed the redetection of the previously annotated 242 snoRNAs included in the Ensembl (v104) database. Our results show that the three algorithms could detect previously annotated snoRNAs with varying efficiency, and while *CMsearch* was able to identify all downloaded sequences as snoRNAs, the efficacy of *snoGPS* and *snoReport* was considerably lower (Supplementary Figure 1). Our redetection experiment suggested that the concomitant use of the three algorithms will result in reduced sensitivity. Yet, as the functional testing of snoRNAs was beyond the scope of this work, we still opted for the most rigorous and conservative approach, and considered only those new sequences in our downstream analyses that were identified by all three *in silico* tools as snoRNAs.

To map the full snoRNAome of zebrafish we first prepared and sequenced brain, liver and gut total RNA samples from two adult males and two adult females, respectively. Our protocol uses small RNA-seq data that are size-selected to capture only RNAs of the required length. During the preparation of our sequencing libraries we paid special attention to size selection and sequenced only the 60-300 bp range. As this size range also encompasses 5S and 5.8S rRNAs (∼120 bp and ∼160 bp, respectively), we computationally removed reads that mapped to rRNA genes from our sequencing datasets before further analysis.

After the elimination of rRNA-specific sequences, we mapped the remaining reads of our small RNA-seq datasets to the zebrafish genome (GRCz11) and created a computational pipeline to identify novel snoRNA genes (Figure 1D). First, we grouped the mapped reads into blockgroups, each blockgroup representing a single ncRNA read profile, which might belong to one of the known ncRNA classes. These ncRNA read profiles are often associated with function (Pundhir et al. 2015), and class-specific discriminative models for C/D box or H/ACA box snoRNAs can be defined. After clustering, if a read profile belonged to one of these ncRNA classes, we considered it a cluster (Figure 1D) (Videm et al. 2014).

The post-mapping analysis of RNA biotypes demonstrated that known snoRNAs were overrepresented amongst the detected RNA species in every tissue type, confirming the validity of our approach (Figure 2A).

**Figure 2:**
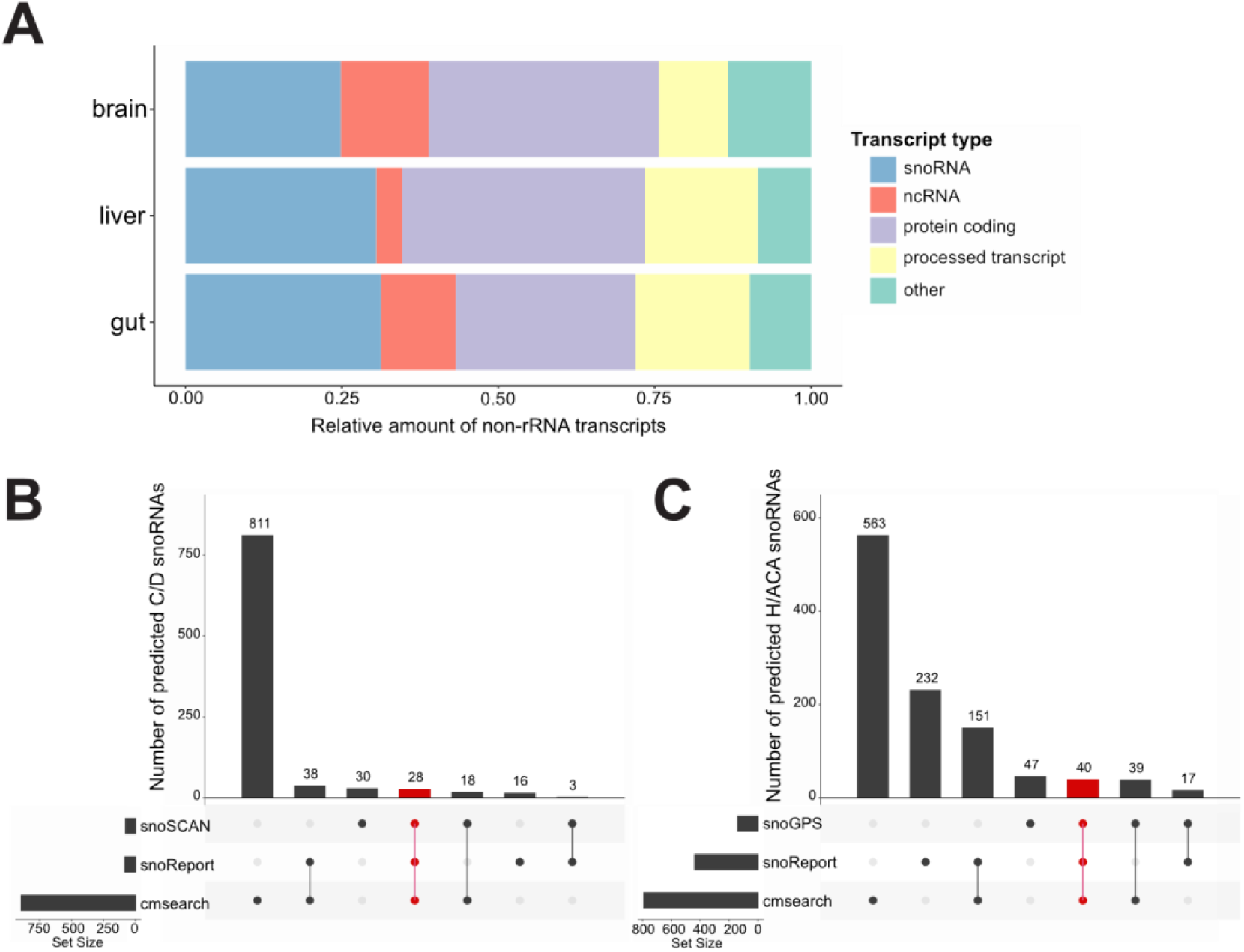
Identification of novel zebrafish snoRNAs. (A) Sequencing the 60-300 nt fractions of zebrafish brain, liver and gut total RNA samples revealed that these are enriched in snoRNAs. (B, C) After removing known snoRNA sequences from our dataset, three different *in silico* prediction methods were used to identify new C/D box (B) and H/ACA box (C) snoRNAs. Up-set plots display the overlap between the individual predictions. (Note: in our analysis we used a conservative approach and accepted as candidates only snoRNAs predicted by all three methods, shown here in red.)

We used our new, tissue-specific expression datasets and combined them with publicly available time series data derived from zebrafish embryos and larvae at different developmental stages (Locati et al. 2017). The presence of several snoRNAs that we had previously repredicted using various snoRNA predictor methods (see above) was also confirmed in these datasets. Off note, this reanalyzed dataset showed comparable abundances for different RNA types as well (Supplementary Table 2).

Using our conservative approach of detection, only considering sequences that were predicted as snoRNAs by all three algorithms used, we were able to identify 67 well-supported new snoRNA-like sequences missing from the current Ensembl database (v104) annotations. Of these novel, putative snoRNAs 27 were C/D box and 39 were H/ACA box and one of them was predicted as belonging to both classes (Figure 2B,C; Supplementary Table 3).

### Characterization of the zebrafish snoRNAs

We compared all our novel, putative snoRNAs with ncRNAs present in RNAcentral (release v17) to identify already annotated zebrafish snoRNAs. This analysis showed that twelve of our novel hits were already present annotated as various kinds of ncRNAs (Supplementary Table 3).

SnoRNAs are most often located in the intronic region of so called snoRNA host genes (SNHGs) but occasionally can also be found in intergenic regions. The expression of intragenic snoRNAs is usually dependent on the expression and/or splicing of their SNHGs, whereas intergenic snoRNAs possess their own promoters (Kufel and Grzechnik 2018; Webster and Ghalei 2023). Most of the previously identified snoRNAs in the zebrafish genome are associated with SNHGs, of which protein coding genes are in excess (Figure 3A). Of the predicted completely novel zebrafish snoRNAs 42 are located within various SNHGs and 13 appear in intergenic positions (Figure 3B). We note that the distribution of known and newly identified genes is highly similar (Figure 3A,B).

**Figure 3:**
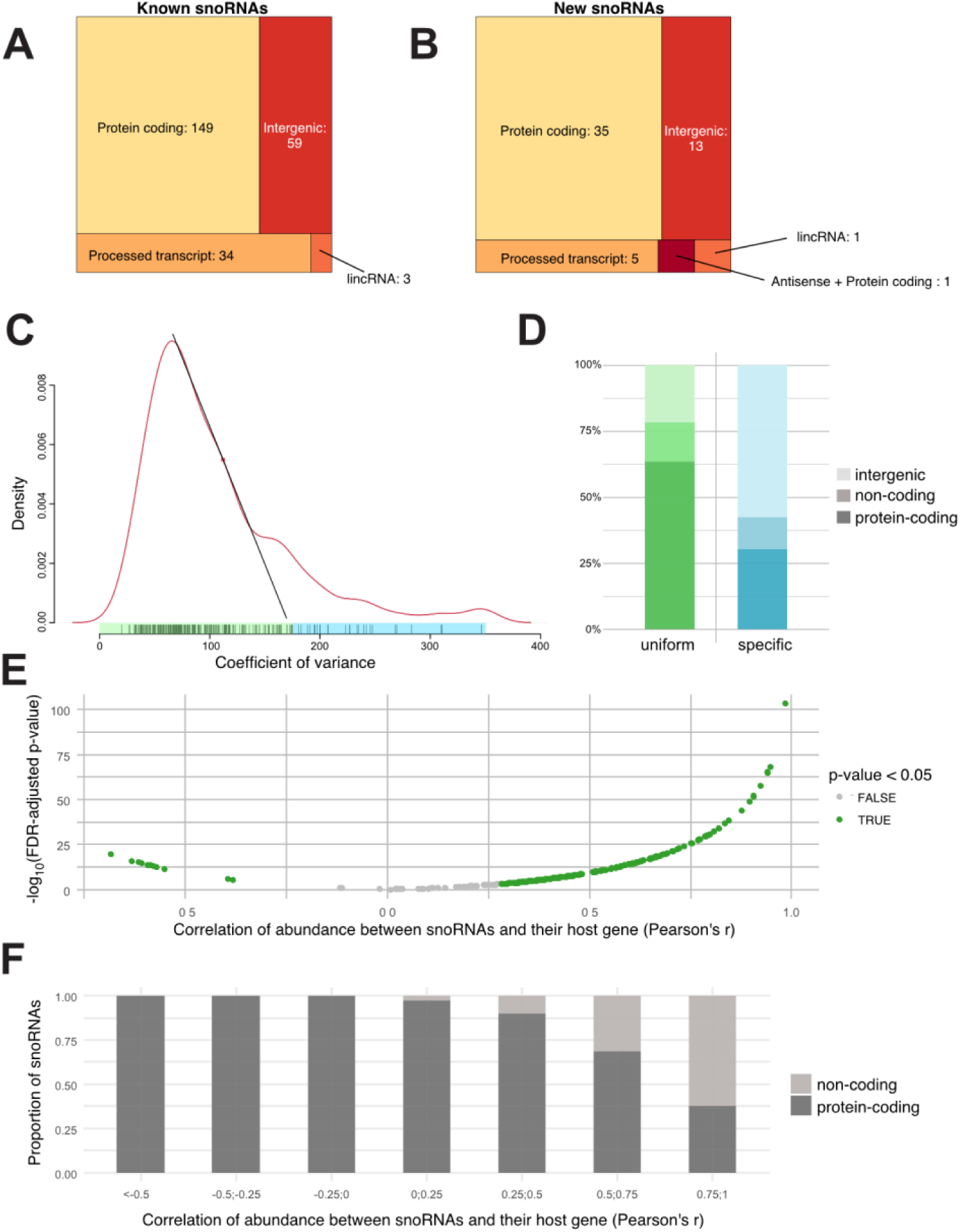
Distribution of novel zebrafish snoRNAs and the characterization of the zebrafish snoRNAome. (A,B) Distribution of known and newly identified snoRNAs shows a similar profile: they are generally associated with host genes (SNHGs), most of which are protein-coding genes. (C) The abundance of zebrafish snoRNAs based on coefficient of variance (CV) values shows a characteristic bimodal distribution. Based on the CV values we can distinguish uniformly expressed (UE) and stage-enriched (SE) snoRNAs (green and blue, respectively). The CV of each snoRNA is indicated as a vertical line on the X axis. (D) The genomic position of UE and SE snoRNAs show differential distribution: the former are enriched in protein coding genes, while the latter in intergenic regions. (E) Correlation between the abundance of snoRNAs and their SNHGs shows that while most often a positive correlation can be detected between the two, for specific snoRNA-SNHG pairs no correlation or outright negative correlation is also possible. (F) SnoRNAs that are positively correlated with their SNHGs can be found in either non-coding or protein-coding SNHGs, whereas lack of correlation or negative correlation is typical for protein coding SNHGs.

To categorize the snoRNAs of the updated zebrafish snoRNAome by their abundance profile we also calculated the coefficient of variation (CV) of their expression (Fafard-Couture et al. 2021). In general, a low computed result for CV corresponds to a more uniform expression across different developmental stages and tissues, typical for housekeeping RNAs, such as transfer RNAs (tRNAs) and snRNAs. In contrast, a high CV corresponds to a more enriched expression in either one or a few stages/tissue types, typical for protein-coding mRNAs and long non-coding RNAs (lncRNAs). Previous analysis of the human snoRNAome already suggested that snoRNA abundance profiles are an intermediate between these two forms, with most snoRNAs showing a uniform expression and a small but significant proportion displaying stage- and/or tissue-specific expression (Fafard-Couture et al. 2021). Our analysis of the zebrafish snoRNAome also suggests that a similar, quasi bimodal CV distribution can be observed (Figure 3C).

The derivative of the CV function also gave us a threshold that we could use to separate uniformly expressed (UE) snoRNAs from those that are stage-enriched (SE) (Figure 3C). Our analysis showed that these two abundance profiles had distinct characteristics. The SE snoRNAs are mostly encoded in the intergenic region, whereas UE snoRNAs are mostly encoded in protein-coding SNHGs (Figure 3D).

As the expression of intron-encoded snoRNAs is dependent on the transcription and splicing of the SNHG, it would be fair to think that the expression level of snoRNA is always correlated positively with that of the host. As it has been shown for human snoRNAs, however, this is not always the case (Fafard-Couture et al. 2021). We calculated the Pearson correlation coefficient (*r*) between the intronic zebrafish snoRNAs and their SNHGs and we could also observe that some snoRNAs have no correlation or even a negative correlation with their SNHG’s expression (Figure 3E). The regulation of SNHG splicing through snoRNAs might be the basis of this phenomenon. Some snoRNAs have been shown to control their SNHGs by promoting non-sense mediated mRNA decay (NMD) (Ojha et al. 2020) and computational analysis shows that several snoRNAs interact with host transcripts, influencing alternative splicing by concealing branch points and shifting the transcript ratio away from NMD (Bergeron et al. 2023). Recent studies provide further insights into the complex relationship between snoRNAs and their SNHGs (Penzo et al. 2020). Of note, in our study, snoRNAs showing no or negative correlation with their SNHGs are almost exclusively found in protein-coding RNAs (Figure 3F). In addition, the expression of long non-coding RNA SNHGs (lncSNHGs) is usually positively correlated with the expression of corresponding snoRNAs (Figure 3F), as reported earlier for lncSNHGs expressed in various immune cells (Xiao et al. 2023).

We also performed gene ontology (GO) analysis, which demonstrated that intragenic snoRNAs that are positively correlated with the expression of their SNHGs show a clear enrichment for processes such as RNA metabolism, ribosome biogenesis or ribosomal protein functions (Supplementary Figure 2).

### Analysis of snoRNA expression patterns during development and in adult tissues

The analysis of the expression dynamics of snoRNAs included in our extended zebrafish snoRNAome demonstrated that some snoRNAs have a dynamic expression during development (Figure 4A), whereas others can have tissue-specific expression patterns in adults (Figure 4B). For example, the expression of one of the newly identified snoRNA genes (*snoRNA-pred2226*) differs strongly between early developmental stages and adult animals (Figure 4A). These dynamic expression patterns suggest that some snoRNAs only function in the early stages of development while others only in adult animals.

**Figure 4:**
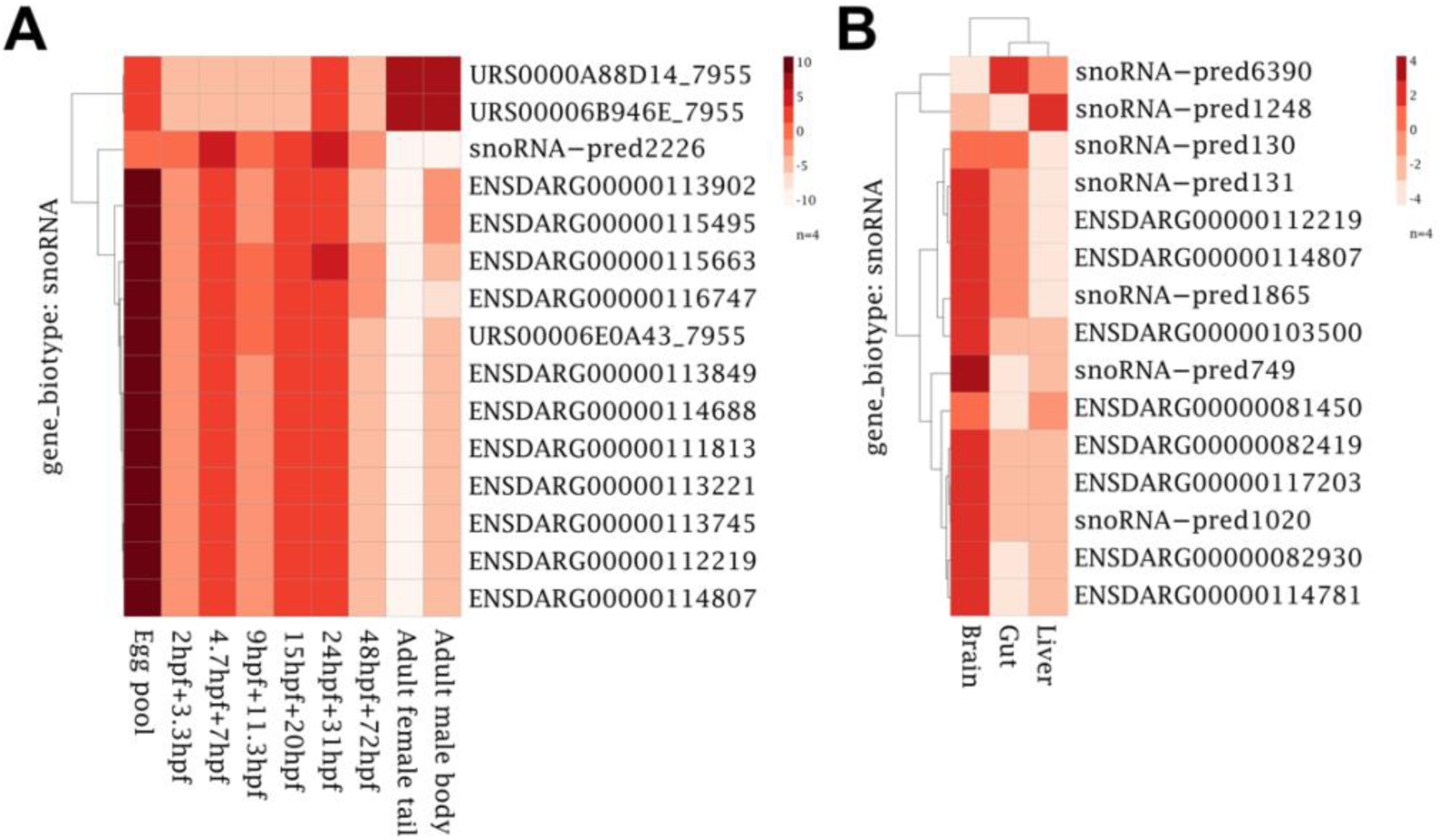
Stage- and organ-specific differential expression of snoRNAs. (A) Some snoRNAs display characteristic stage-specific expression patterns. Whereas some snoRNAs are highly expressed during embryogenesis and at much lower levels in adults, other snoRNA genes show a complementary expression pattern. (B) Adult datasets show differential expression of particular snoRNAs in different organs. Off note, organs with linked (endodermal) developmental origin, such as the gut and the liver, show similar snoRNA expression patterns.

Another seven newly annotated snoRNAs (*snoRNA-pred6390, snoRNA-pred1248, snoRNA-pred130, snoRNA-pred131, snoRNA-pred1865, snoRNA-pred749* and *snoRNA-pred1020*) show strong expression differences in different organs. Their expression in endodermal tissues, such as the liver and the gut is more similar to each other, and differs markedly from that in neuroectoderm.

### snoDanio: The zebrafish snoRNAome database

Using our extended information of the zebrafish snoRNAome we created an interactive database, snoDanio (https://renata-h.shinyapps.io/98665a9405b44ede86eeb7179988104f/), an openly licensed resource that facilitates integrative and interactive display and analysis of various zebrafish snoRNAs by mining multiple high-throughput sequencing data from tissue and developmental stages of zebrafish (Figure 5).

**Figure 5:**
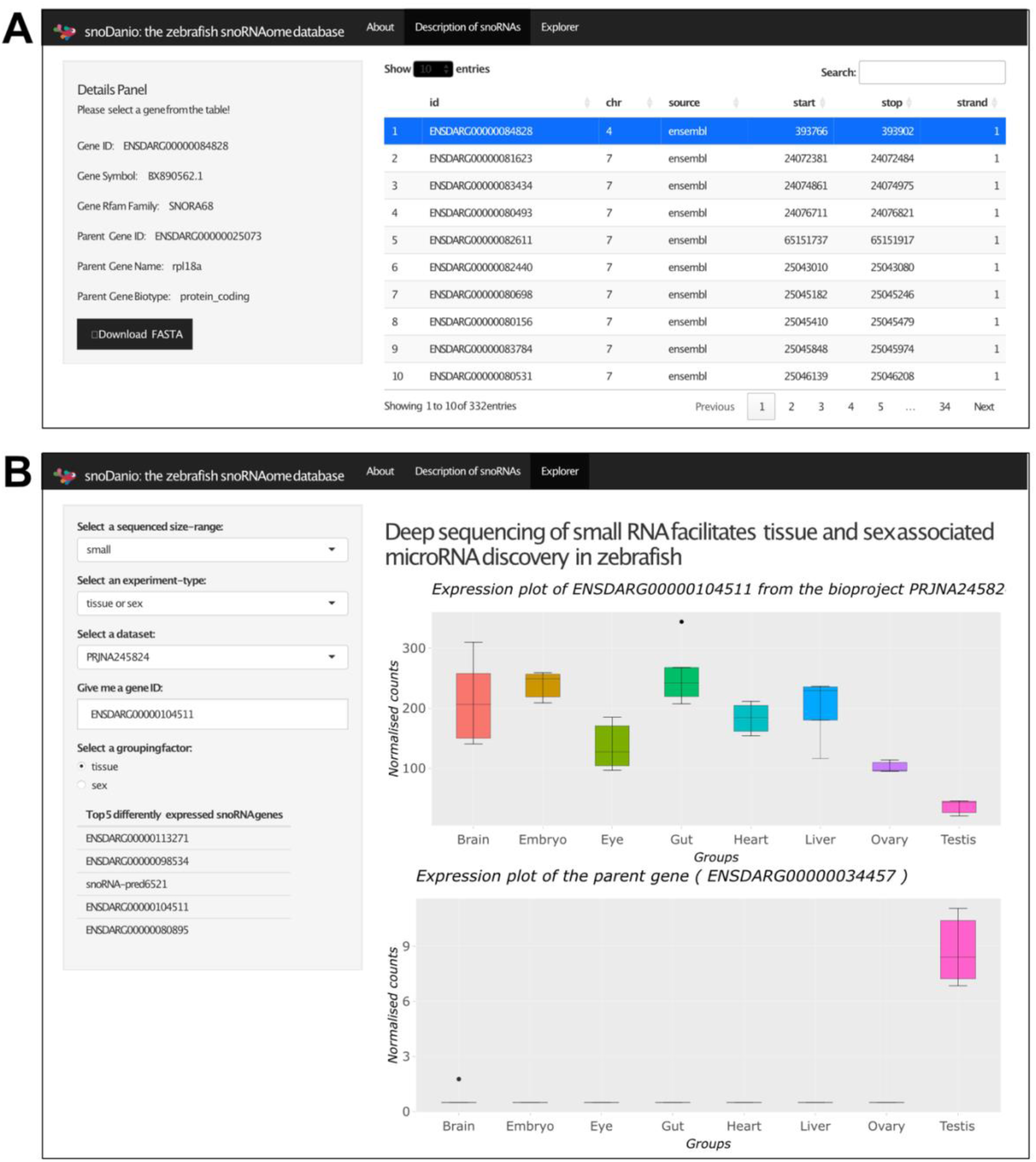
Page design for snoDanio. (A) Basic description of snoRNAs including genomic location, snoRNA classification and SNHG information. (B) Expression analysis of respective snoRNAs and SNHGs using different datasets. Sequencing data are categorized by size-range and experiment type. (Note that grouping factor will depend on the design of the original sequencing experiment.)

Users can access basic, descriptive information of individual snoRNAs, including their classification, genomic position and SNHG information (Figure 5A), but can also assay the expression of particular snoRNAs in the preprocessed datasets already included in the database (Figure 5B). These preprocessed datasets are categorized on the basis of the size-range of RNAs included in the sequencing (mid, polyA, polyA-depleted, small, total, and total or polyA), and on the nature of the experiment design (assessing tissue-specific, sex-specific, developmental/ time-related or treatment-related changes), as determined from the metadata of the uploaded datasets. Depending on the type of the original experiment, multiple groupings for visualizations are also possible. The expression results and the sequence of snoRNAs can be downloaded on demand in image and FASTA formats, respectively, for further use.

## DISCUSSION

The expanding repertoire of putative and documented snoRNA functions (McMahon et al. 2015; Deogharia and Majumder 2018; Liang et al. 2019; Bergeron et al. 2020; Huang et al. 2022) suggests that the significance of this ncRNA class has been hitherto underappreciated, and new insights into the regulation of gene expression will be gained by the in-depth study of these molecules. It is important, therefore, to compile comprehensive lists of snoRNAs and to document their dynamic and tissue-variable expression in all model organisms.

In the current study we set out to define a well-supported list of snoRNAs expressed in zebrafish during development and in adults and create an interactive database to analyse the data. Using size-fractionated (60-200 bp) sequencing datasets, rigorous filtering pipelines and relying on predictions by multiple annotation algorithms (*snoReport*, *snoGPS* and *CMsearch*) we achieved identification of 67 novel snoRNA genes with high confidence (Figure 2B,C). Some of these genes are situated in intergenic regions, but the majority of them are encoded within the introns of protein-coding and lncRNA host genes. With our newly discovered snoRNAs, the complete, annotated zebrafish snoRNAome stands at 435 genes. We note that this figure is likely a conservative estimate as we only considered genes that passed our stringent filtering pipeline (Figure 1D). Therefore, by sequencing further tissues and by employing better annotation tools identification of further zebrafish snoRNAs is likely.

Our new dataset allowed us to compare the composition and expression dynamics of the human and zebrafish snoRNAomes. We observed that while most of the snoRNAs are uniformly expressed in both species, a subset of them shows tissue-specific expression.

To date, multiple mechanisms have been identified that explain the effect of snoRNAs on gene expression. The most abundant target of snoRNA modification is rRNA and the role of snoRNA-guided modifications in ribosomogenesis and translational fidelity is well documented (Liang et al. 2009; Jack et al. 2011; Watkins and Bohnsack 2011; Sloan et al. 2016; Ojha et al. 2020; Zhao et al. 2022; Dörner et al. 2023; Zhao et al. 2023). Defective ribosomal pseudouridylation and 2’-*O*-methylation can lead to developmental defects (He et al. 2002; Ruggero et al. 2003; Ge et al. 2010; Pereboom et al. 2011; Zhang et al. 2012; Anchelin et al. 2013; Bellodi et al. 2013; Delhermite et al. 2022; Breznak et al. 2023; Ni and Buszczak 2023) and various diseases, including cancer (Bellodi et al. 2010; McMahon et al. 2015; Bohnsack and Bohnsack 2019; Balogh et al. 2020; Kampen et al. 2020; Kang et al. 2021; Barozzi et al. 2023). Multiple studies have also demonstrated that snoRNAs themselves can be effectively used as biomarkers for numerous malignancies (Dong et al. 2008; Zhou et al. 2017; Deogharia and Majumder 2018; McMahon et al. 2019; Huang et al. 2022; Zhang et al. 2022) and recently they have been also linked to cellular senescence (Cheng et al. 2024).

Accumulating evidence suggests that pseudouridylation of mRNAs is also abundant and can affect the efficiency of gene expression (Carlile et al. 2014; Schwartz et al. 2014; Li et al. 2015; Martinez et al. 2022; Nir et al. 2022; Rodell et al. 2023; Zhang et al. 2023; Sun et al. 2023; Hamar and Varga 2023). As different studies identified different roles for pseudouridines in mRNA (Dai et al. 2022; Pederiva et al. 2023), the exact role for this modification (which is most likely to be context-dependent) remains to be confirmed.

Alas, cell-type specific presence of particular posttranscriptional modifications in rRNA has been also described and linked to ribosomal heterogeneity in a variety of organisms (Chikne et al. 2016; Hebras et al. 2020; Marchand et al. 2020; Rajan et al. 2023). It has been proposed that differential epitranscriptomic modification of particular rRNA nucleotides could result in specialized ribosomes that are more adept in translating the messages in the given tissue-type(s) (Xue and Barna 2012; Shi and Barna 2014; Sloan et al. 2016; Ni and Buszczak 2023). Off note, it has been already demonstrated that knock-down of the U26, U78 and U44 C/D-box snoRNAs (driving methylation in 28S and 18S rRNAs, respectively), or depletion of U8 (which is required for removal of the 3′ external transcribed spacer or ETS sequence) can yield in divergent developmental phenotypes in zebrafish embryos (Higa-Nakamine et al. 2012; Badrock et al. 2020).

Our update of the zebrafish snoRNAome has also indentified multiple snoRNAs with sex-, stage- and tissue-specific expression, suggesting a hitherto underappreciated dimension of gene regulation. Understanding how this diversity in snoRNA expression can contribute to the developmental process and modulate the function of other genes should be the focus of later studies. Indeed, bisulfite-induced deletion sequencing (BID-seq) analysis of different mouse tissues has already uncovered a high level of differential pseudouridylation in the transcriptome of different organs in mice (Dai et al. 2022), and differential maternal and zygotic transcription of certain zebrafish snoRNAs has been also described before (Pagano et al. 2019).

The epitranscriptomic methodological revolution brought into focus the importance of several post-transcriptional modifications in gene expression regulation and also made it abundantly clear that these modifications can greatly vary between tissues, developmental stages and physiological conditions (Hamar and Varga 2023). The snoRNA-guided function of the fibrillarin- and dyskerin-containing RNPs makes them ideally suited to have central roles in the dynamic regulation of the epitranscriptome and through their effect on ribosome biogenesis, on the translational process as well. Understanding, therefore, how the function of these two essential enzymatic complexes can be regulated by the presence or absence of particular snoRNA guides will be a central question in this field in the coming years.

Our novel database, which compiles multiple lines of evidence to provide a comprehensive expression atlas for the snoRNAome of one of the most versatile genetic model organisms will aid this work by helping to generate novel hypotheses and to identify essential snoRNAs for particular biological processes. We plan to regularly update the snoDanio database by performing the reanalysis of newly available datasets. This approach will give us a chance to discover and characterize further snoRNAs. Moreover, if robust target prediction and validation pipelines can be developed, our data could be also integrated into more specific signaling network databases, such as SignaLink (which already handles zebrafish data) (Csabai et al. 2021) or the Zebrafish Information Network (ZFIN) (Bradford et al. 2022) as well.

## Data availability

The new sequencing data generated during this project can be found at the NCBI SRA archive PRJNA1127032 (reviewer link: https://dataview.ncbi.nlm.nih.gov/object/PRJNA1127032?reviewer=so7fbbcj3re2ptlll0am6dd8nj). We have also reused and reanalyzed data from the PRJNA347637 BioProject (Locati et al. 2017). Further data related to this research project is available at Github (https://github.com/danio-elte/Danio_snoRNAome) and Zenodo (Hamar and Varga 2024).

## Acknowledgments

We would like to thank Anita Rácz for fish care and Julianna Víg for help with the proofreading. This work was supported by the National Research, Development and Innovation Fund of Hungary grants FK124230 and K146460, the ELTE Eötvös Loránd University Institutional Excellence Program Grant 1783-3/2018/FEKUTSRAT and the Central European Leuven Strategic Alliance grant CELSA/22/026. MV is a János Bolyai fellow of the Hungarian Academy of Sciences and he is also supported by the ÚNKP-23-5 New National Excellence Program of the Ministry of Culture and Innovation from the source of the National Research, Development and Innovation Fund. The authors also acknowledge the support of the Freiburg Galaxy Team: Björn Grüning, Bioinformatics, University of Freiburg (Germany) funded by the German Federal Ministry of Education and Research BMBF grant 031 A538A de.NBI-RBC and the Ministry of Science, Research and the Arts Baden-Württemberg (MWK) within the framework of LIBIS/de.NBI Freiburg.

## Author contributions

Conceptualization: RH, MV.

Data curation: RH.

Funding acquisition: MV.

Investigation: RH, MV.

Methodology: RH.

Project administration and supervision: MV.

Writing – original and revised text: RH, MV.

