## Supplementary Figures for "The zebrafish (*Danio rerio*) snoRNAome"

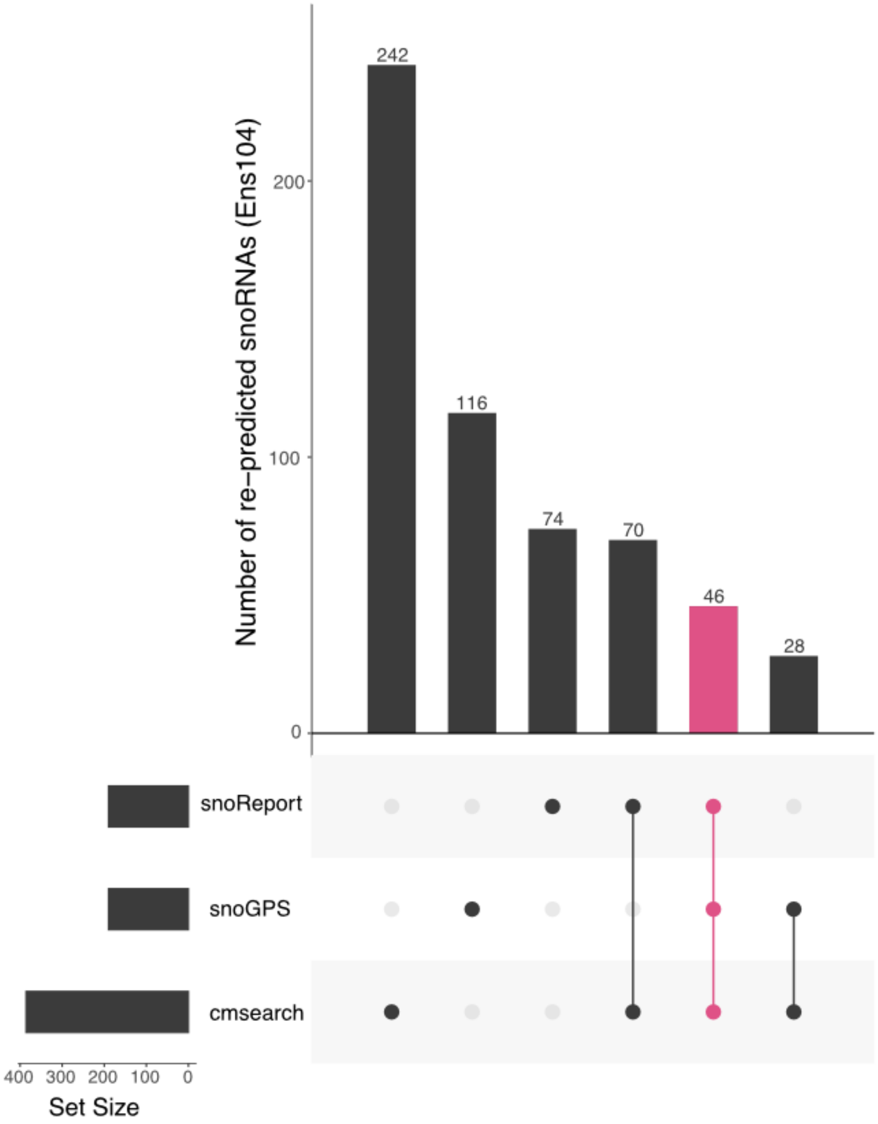


Supplementary Figure 1: Redetection of snoRNAs previously annotated in Ensembl. Three algorithms – *cmsearch*, *snoReport* and *snoGPS* – were tested, to estimate their efficiency in redetecting previously annotated snoRNAs. Of these only *cmsearch* was able to detect all 242 snoRNAs.

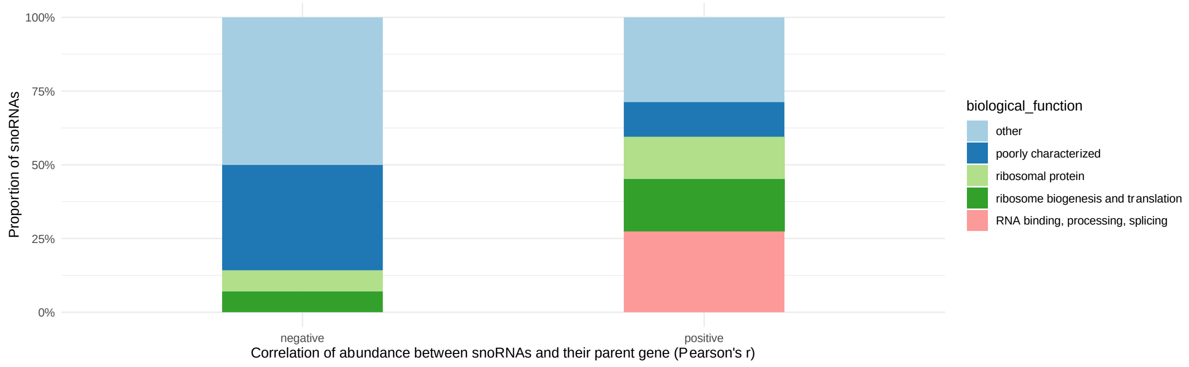


Supplementary Figure 2: GO ontology analysis of SNHGs which show either a negative (left) or a positive (right) association with the hosted snoRNA.
